# Regulation of excitatory presynaptic activity by Ambra1 protein determines neuronal networks in sex-dimorphic manner

**DOI:** 10.1101/2022.12.19.521004

**Authors:** Anes Ju, Bekir Altas, Hong Jun Rhee, Manuela Schwark, Imam Hassouna, Albrecht Sigler, Hiroshi Kawabe, Kamal Chowdhury, Hannelore Ehrenreich, Nils Brose, JeongSeop Rhee

**Author notes:** Samsung Advanced Institute of Technology, Samsung Electronics, Suwon-si, South Korea (Anes Ju).

## Abstract

Heterozygous mutation of *Ambra1*, known as a positive autophagy regulator, produces autism-like behavior in mice and autistic phenotypes in humans in a female-specific manner. However, the substantial roles of the *Ambra1* mutation in neurons are still unknown. We find that *Ambra1* heterozygotes display a moderate decrease in excitatory synaptic release *in-vitro* and *ex-vivo* exclusively in females without autophagy activity, resulting in significant alterations in γ-oscillation power and seizure susceptibility by excitatory/inhibitory (E/I) imbalance. Specifically, *Ambra1* deficiency has no effect on neurogenesis and morphogenesis, but selectively decreases excitatory synaptic activity without changes in synapse number, quantal size, synaptic release probability, and synaptic plasticity. Therefore, the limited excitatory synaptopathy by Ambra1 expression levels ultimately determines E/I imbalance in global neural networks leading to the female-specific ASD.

## Introduction

Autism-spectrum disorder (ASD) is a neurodevelopmental disorder mainly characterized by deficits in social interaction/communication and restricted/repetitive patterns of behavior (Association, 2013). Epidemiological studies estimated that ASD has been diagnosed in more than 1% of the world’s population (Elsabbagh et al., 2012; Vos et al., 2016) and described as a sexually-dimorphic disease, with four times more males than females being diagnosed (Baron-Cohen et al., 2011; Chakrabarti and Fombonne, 2001). Genetic etiology of ASD is highly heterogenous, with greater than 100 identified risk genes involved in diverse functions, such as transcriptional regulation, protein synthesis and degradation, synapse function and synaptic plasticity (Bourgeron, 2015; Delorme et al., 2013; Ebert and Greenberg, 2013). Whether genetically distinct forms of ASD share common pathophysiology at the neural network level still remains to be elucidated.

Emerging evidence has suggested a disturbed homeostasis of excitatory/inhibitory (E/I) balance as an etiology of ASD (Nelson and Valakh, 2015; Zikopoulos and Barbas, 2013). Epilepsy, relatively high comorbidity in ASD, occurring in 5-38% of autistic individuals, highlights the possibility of shared neurophysiological mechanisms involved with changed neural network activities (“Epilepsy and autism spectrum disorders may have a shared aetiology,” 2016; Lee et al., 2015). This hypothesis is corroborated by electroencephalogram (EEG) abnormalities often observed in patients with ASD or epilepsy (Martinerie et al., 1998; Mathalon et al., 2015; Rossi et al., 1995; Spence and Schneider, 2009), addressing that altered neuronal network and synchronicity are related with those two diseases. Based on several mouse studies proposing sexually dimorphic mechanisms regulating neural circuits (Li et al., 2016; Malishkevich et al., 2015), E/I balance is a critical factor of ASD in a sexually-dimorphic manner. However, the neurophysiological substrates of E/I imbalance in ASD still stimulates our curiosity.

Ambra1 (activating molecule in Beclin1-regulated autophagy) is a crucial regulator in autophagy, proliferation and apoptosis in eukaryotic cells (Maria Fimia et al., 2007). Homozygous mutation of *Ambra1* gene in mice (*Ambra1^gt/gt^*) resulted in embryonic lethality showing neural tube defects (Maria Fimia et al., 2007). Interestingly, *Ambra1* heterozygous mice (*Ambra1^+/gt^*), that are viable, produced clear autism-like behaviors only in females, which might be linked to sexually-dimorphic expression of Ambra1 protein in brain tissue (Dere et al., 2014). Additionally, our previous study showed early brain enlargement and different seizure propensity depending on developmental stages in this mouse line in a female-specific manner supporting *Ambra1^+/gt^* mice as a model of female-specific ASD (Mitjans et al., 2017). Especially, this study includes human genetic research reporting a significant association between autistic features and intronic single nucleotide polymorphisms of the *AMBRA1* gene in females but not in males. Therefore, *Ambra1* heterozygous mice is proved as a construct-valid genetic mouse model of female ASD (Mitjans et al., 2017).

The present study has been designed to explore the neural substrate underlying this E/I balance observed in the brains of *Ambra1^+/gt^* females by screening functional and morphological aspects of neural network in brain slice. In addition, in order to define the precise role of the Ambra1 protein, we studied the functional consequences by the absence of *Ambra1* gene in autaptic neuronal culture of *Ambra1* homozygous mutation. Our study revealed that Ambra1 present in the brain was limitedly located in neuronal cells, and was particularly accumulated in the synapses. We found that, regardless of sex, Ambra1 was directly involved in excitatory synaptic activity, while showing no effect on neuronal development and synapse formation. More importantly, it was noticed that the reduction of excitatory synaptic activity by *Ambra1* heterozygosity only in females, although not significant, is a decisive cause of synaptic E/I input imbalance, contributing to ASD.

## Results

### Region-, cell type- and subcellular-specific expression of Ambra1 protein

We first analyzed the expression pattern of Ambra1 protein in region, different cell types and subcellular location of mouse brain. mRNA expression data from Allen brain atlas revealed that *Ambra1* is widely present (Figure 1A). Histochemical or immunofluorescent staining of β-galactosidase (β-gal) in mouse brains showed that Ambra1 is abundantly expressed in cortex, striatum and hippocampus and in neurons, but not in glial cells (Figure 1B-E). Subcellular fractionation of *Ambra1^+/+^* cortex (Figure 1F) (Bermejo et al., 2014), where the purification of synaptic membrane was validated (Figure 1G), was used for Western Blot of Ambra1 protein. Surprisingly, Ambra1 protein was identified not only in ER-Golgi enriched fractions (P2B) but also in crude synaptic membrane (CSM) and pure synaptic membrane fractions (SM, Figure 1G). These data illustrate that Ambra1 protein is located only in neurons and is particularly distributed in synapses, suggesting a possibility of its role in neuronal communication.

**Figure 1:**
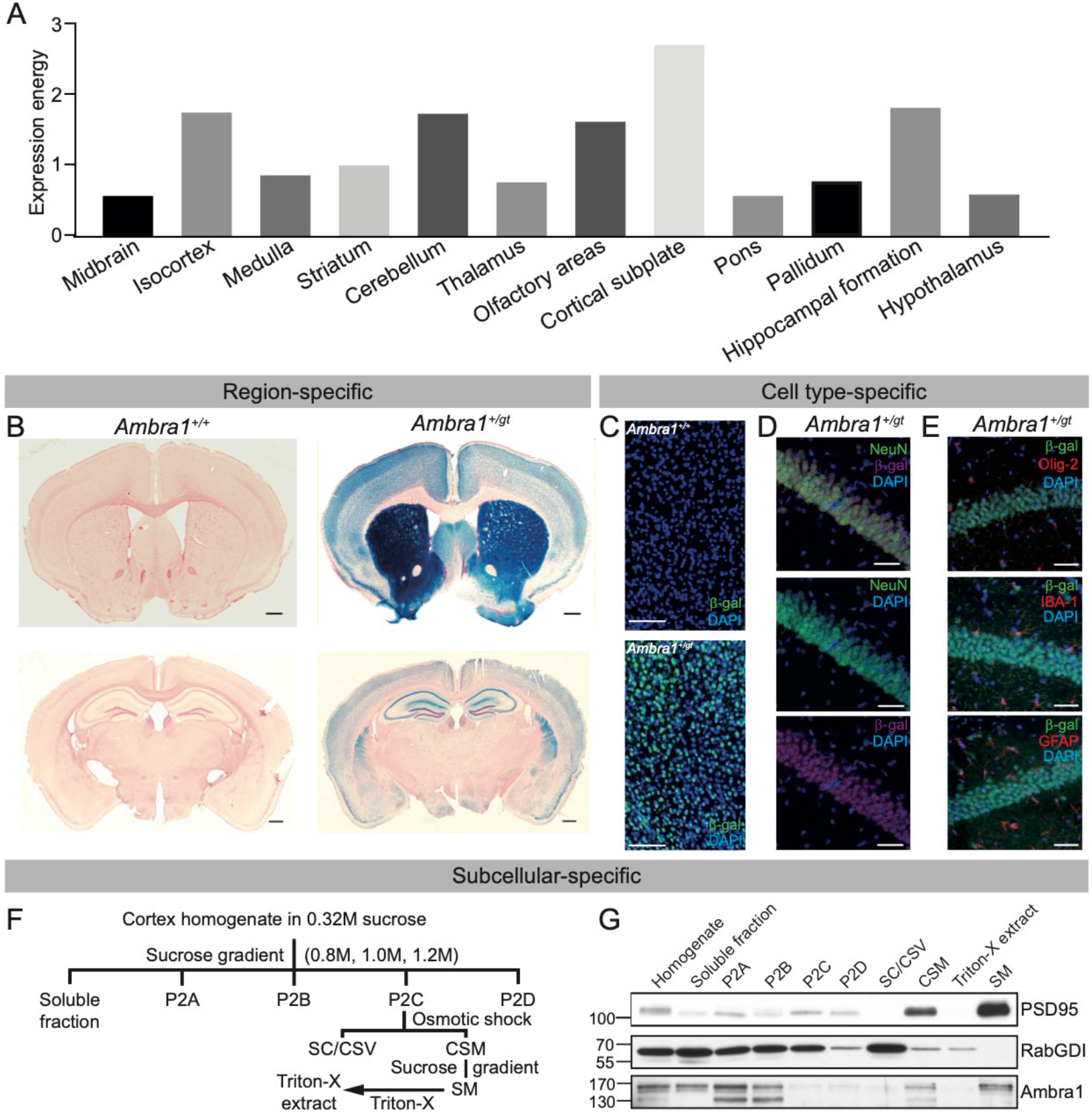
Region-, cell type- and subcellular specific location of Ambra1 protein in the brain. **A** Expression level of Ambra1 mRNA in different brain regions of a mouse (n=1). **B** X-galactosidase staining in coronal brain section of *Ambra1^+/+^* and *Ambra1^+/gt^* mice. Scale bar, 100 μm. **C** Immunofluorescence staining of β-galactosidase (β-gal). **D, E** Co-staining of β-gal with different cellular markers, including neuronal marker, NeuN (D) or glial markers, Olig-2, IBA-1 and GFAP (**E**) in hippocampal CA1 pyramidal region of *Ambra1^+/gt^* mouse. Scale bar, 40 μm. **F** Schematic representation of the subcellular fractionation step. P2A, Myelin-enriched fraction; P2B, ER/Golgi-enriched fraction; P2C, Synaptosome fraction; P2D, Nucleus- and mitochondria-enriched fraction; SC, Synaptic cytoplasm; CSV, Crude synaptic vesicles; CSM, Crude synaptic membrane; SM, Pure Synaptic Membrane Fractions. **G** Western blots of PSD95, RabGDI and Ambra1 proteins in subcellular fractions of *Ambra1^+/+^* mouse cerebral cortex. PSD95 and RabGDI were used for verification of purification of SM fraction.

### No change in activity-dependent synaptic plasticity upon Ambra1 heterozygous mutation

Due to neuronal expression and synaptic location of Ambra1 protein, we sought to determine the consequences of *Ambra1* heterozygous mutation in neural networks for learning and memory. Based on previous studies showing the modification of synaptic plasticity and oscillatory activity in several ASD mouse models (Hammer et al., 2015; Mathalon et al., 2015), we recorded them in acute hippocampal slices from 4 week-old mice before sexual maturation (Heiniger H. J. and Dorey, 1989), using extracellular recording. Overall, we found the input-output curve, paired-pulse ratio, and early-phase long-term potentiation (Figure 2) were comparable between two genotypes in male and female mice, pointing out that, upon *Ambra1* heterozygous mutation, activity-dependent synaptic activities and plasticity are unaltered.

**Figure 2:**
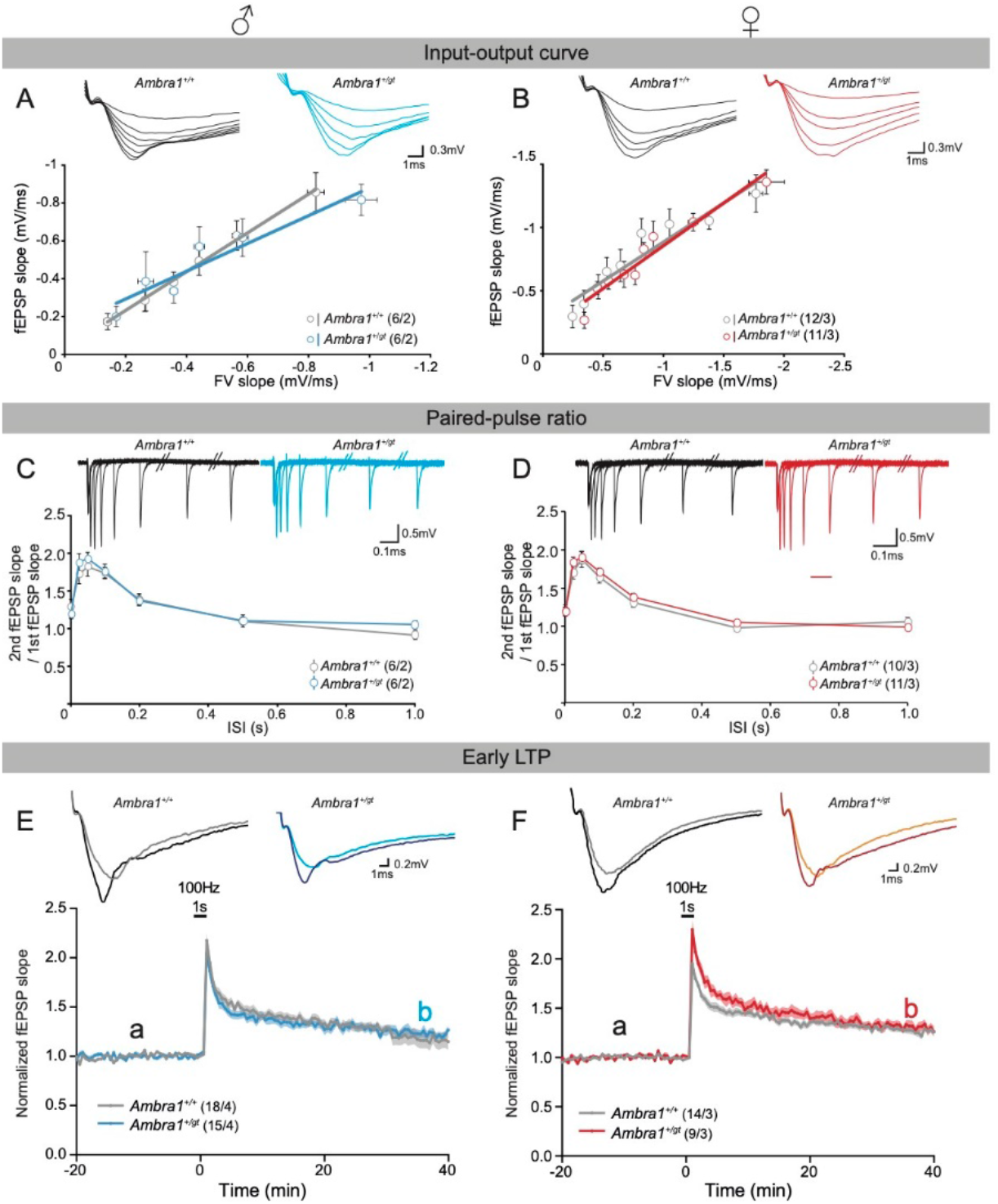
Activity-dependent synaptic transmission and short- and long-term synaptic plasticity of hippocampal CA1. Left panel contains data from male mice, while right panel represents data from female mice. Field excitatory postsynaptic potential (fEPSP) slope was recorded in the *Striatum Radiatum* of CA1 upon Schaffer-Collateral stimulation in acute hippocampal slice. **A, B** Input-output curve from *Ambra1^+/+^* (black) and *Ambra1^+/gt^* (color) mice in both sexes. fEPSP slopes were measured after increasing the stimulus intensity and plotted along their fiber volley (FV) slopes. **C, D** Paired-pulse ratio curve from two genotypes in both sexes. Ratios of 2^nd^ and 1^st^ fEPSP slope (Paired-pulse ratio) after two stimuli within different time intervals (Interstimulus interval, ISI) were plotted along their respective intervals. Representative traces were shown within graphs. **E, F** Early-phase long-term potentiation (E-LTP) from two genotypes in both sexes. fEPSP slopes after stimuli every 30 second were normalized to baseline and plotted along time. After 20 minutes of baseline, a high frequency stimulation (100 Hz for 1 sec) was given to induce potentiation. Representative traces are shown within graphs (light color: baseline, dark color: potentiation). N numbers are written next to the legend within the graphs and shown as slice number/animal number. Mean ± S.E.M. are presented in line and area and statistical difference was defined by p-value between genotypes from Repeated-Measures of ANOVA.

### Perturbed γ-power and seizure propensity by Ambra1 heterozygous mutation only in females but not in males

To specifically and concretely measure the local activity of neural network, we assessed the oscillatory activity in hippocampal CA3 pyramidal layer induced by Kainate. The peak frequencies were detected within γ-range (25-45 Hz) and comparable between two genotypes. Intriguingly, the average power of γ-oscillation was significantly lower in *Ambra1^+/gt^* females compared to control littermates, while male mice exhibited similar levels between two genotypes (Figure 3D-E), indicating female-specific alteration of synchrony of neural network activities in *Ambra1^+/gt^* brains.

**Figure 3:**
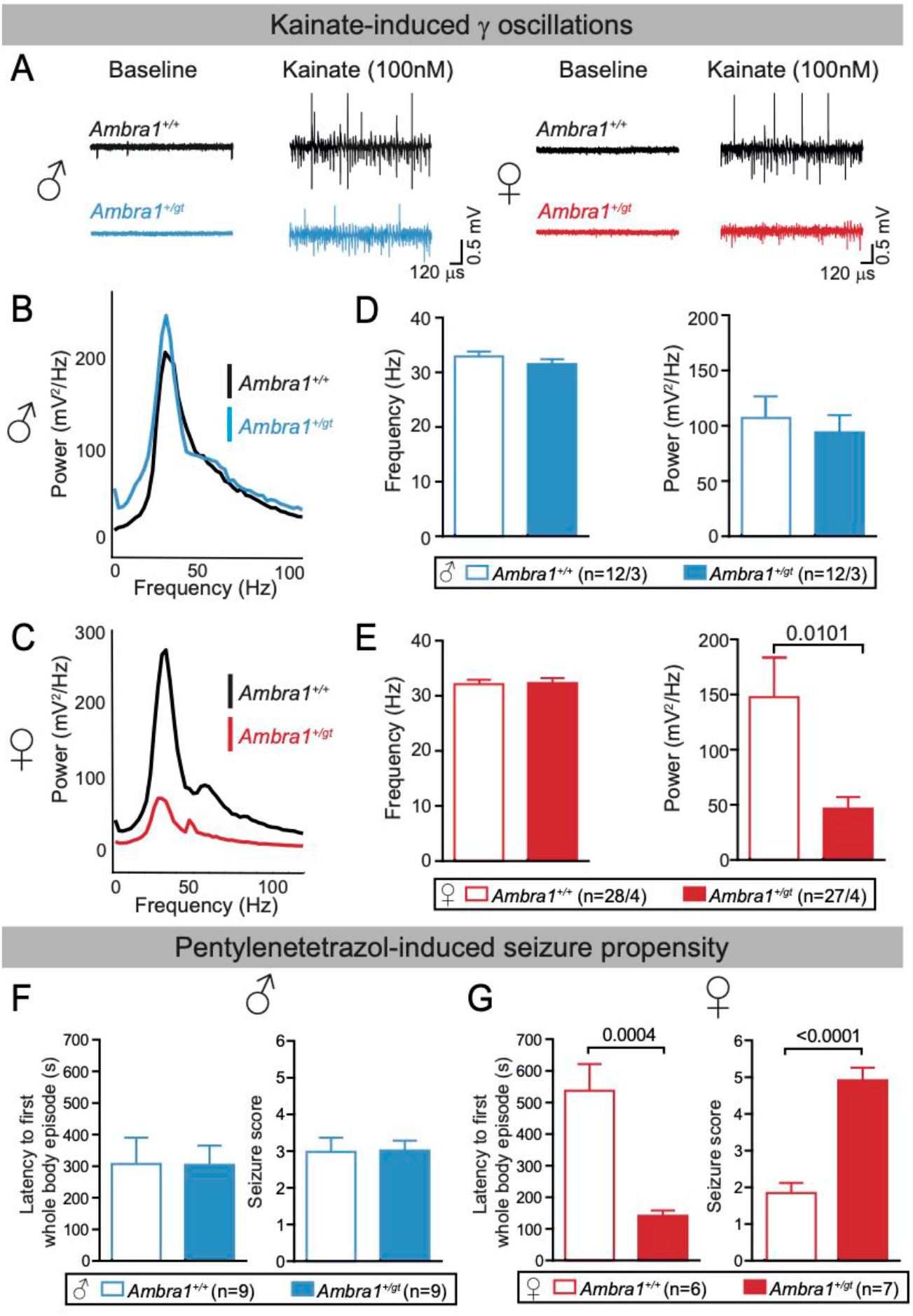
Perturbed γ-power and seizure propensity only in *Ambra1^+/gt^* females but not in males. **A** Representative traces of **γ**-oscillations before and during being induced by 100 nM kainate in hippocampal CA3 pyramidal layer of acute brain slices in *Ambra1^+/+^* and *Ambra1^+/gt^* male and female mice. **B, C** Representative power spectrums of gamma oscillations are shown. **D, E** Quantifications of the frequency of maximum power and average power within gamma range (25-45 Hz) at 4 weeks old. **F, G** Latency to first whole-body episode and seizure score were measured during observation after injection of pentylenetetrazol (50mg/kg) in 12-13 weeks old mice. The bar graphs are presented as mean ± S.E.M and slice number/animal number or animal numbers of each group are noted next to the legends. Statistical analysis between *Ambra1^+/+^* and *Ambra1^+/gt^* was performed by two-tailed unpaired t-test with significance level p < 0.05.

Epilepsy, one of the comorbid conditions of ASD (Bolton et al., 2011), is a behavioral feature manifested by abnormal synchrony of neural network activities (Sun et al., 2021). The seizure threshold was markedly higher in *Ambra1^+/gt^* females compared to control, which is shown by earlier latency of whole-body seizure episodes and higher seizure score, whereas those parameters were similar between two genotypes in males (Figure 3F-G). Taken together, higher seizure propensity and lower power of gamma oscillations in heterozygous females demonstrate that female mice are more sensitive to disturbed synchronous network activity upon *Ambra1* heterozygous mutation.

### E/I imbalance by change in functional excitatory synapses, regardless of autophagy activity

To investigate the cellular substrates underlying the E/I imbalance which is the significant mechanism causing the abnormal synchronization in a neuronal network, we first compared the population of excitatory and inhibitory neurons between two genotypes. The densities, sum and ratio of the mature glutamatergic (CTIP2+) and GABAergic (GAD67+) neurons as well as the density of parvalbumin-expressing (PV+) interneurons were similar in pyramidal layer of hippocampus between *Ambra1^+/+^* and *Ambra1^+/gt^* female mice (Figure 4A-G). This data suggests that the *Ambra1* heterozygous mutation is not crucial for neuronal proliferation and apoptosis, which are not a main factor for the alteration in γ-oscillation power in this mouse line.

**Figure 4:**
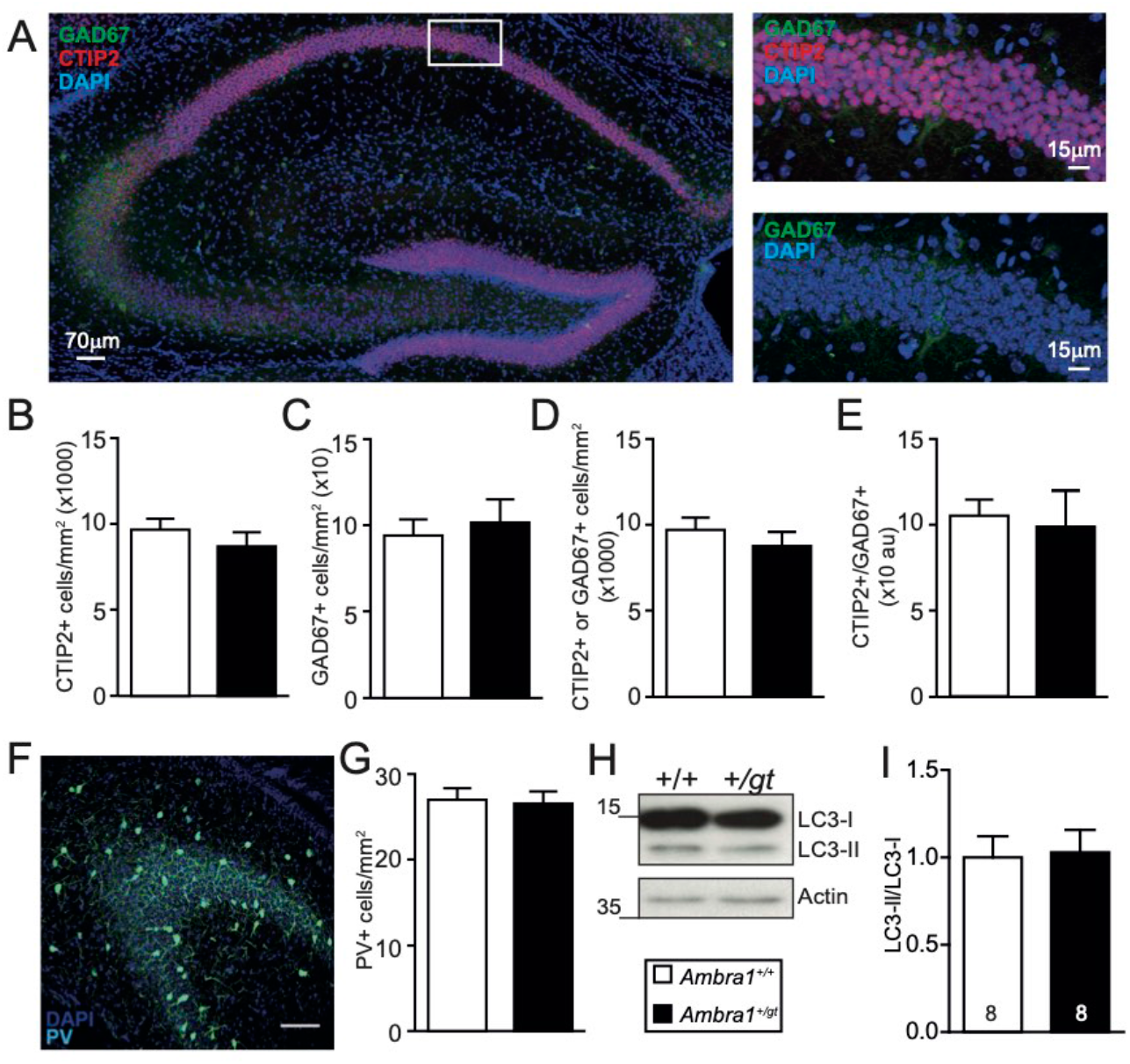
Comparable neuronal numbers and autophagic activity in the brains of *Ambra1^+I+^* and *Ambra1^+Igi^* mice. **A** Example images of hippocampal CA1 region immunostained with anti-CTIP2 and anti-GAD67 antibodies. **B-E** The quantifications of the density of CTIP2+ (**B**), GAD67+ (**C**), sum of CTIP2+ and GAD67+ neurons (**D**) and ratio of CTIP2+ and GAD67+ neurons (**E**) in *Ambra1^+/+^* and *Ambra1^+/gt^* mouse hippocampus. **F** Example images of hippocampal CA3 region immunostained with an anti-PV antibody. 4 weeks old wild type mouse was used. **G** The quantifications of PV+ neurons in *Ambra1^+/+^* and *Ambra1^+/gt^* mouse hippocampus. 13-15 sections in 1-2 animals were used for analysis per group. **H** Sample picture of LC3 Western Blot between cortical homogenates of both genotypes (n=8 for each group) in female mice. The data was normalized to the average value of wild-type group. **I** Comparison of LC3-II and LC3-I intensities between two genotypes. The bar graphs represent mean ± S.E.M and statistical analysis was performed by two-tailed unpaired t-test with significance level p < 0.05.

By Western blotting with the anti-LC3 antibody, LC3-II/LC3-I ratio, an indicative of autophagic activity, was similar between *Ambra1^+/+^* and *Ambra1^+/gt^* in female cortical homogenates (Figure 4H-I). This is corroborated by unaltered expression levels of different synaptic proteins in hippocampal homogenates or cortical synaptosomal fractions between two genotypes (Figure S1). Therefore, our data imply that the autophagic activity of Ambra1 unaccompanied by neuron is not a critical factor in the E/I imbalance.

We measure miniature excitatory and inhibitory postsynaptic currents (mEPSC and mIPSC) at the same neuron in acute brain slice and morphological features by simultaneously filling the recorded neurons with biocytin, as a minimal functional model system (Figure 3A).

The morphological properties of hippocampal pyramidal neurons, including dendritic arborization and number of mushroom spines (Figure 5A-C), were unaltered by *Ambra1* heterozygous mutation, which is additionally supported by independent experiments using *in utero* electroporated samples (Figure S2). Moreover, PSD95 and Gephyrin as well as PSD95/Gephyrin expression, implying the number of excitatory and inhibitory post-synapses and their ratio, were not altered in *Ambra1^+/gt^* brains (Figure S1C-D). Similar neuronal morphology and expression levels of postsynaptic proteins can infer that the number of excitatory and inhibitory postsynapses were unchanged by *Ambra1* heterozygous mutation. Without changes in mEPSC and mIPSC amplitudes, only mEPSC frequencies of female *Ambra1^+/gt^* neurons showed a strong tendency for reduction (Figure 5D-F), suggesting that the possibility due to a decrease in the number of functional glutamatergic synapses cannot be excluded. Surprisingly, the ratio of frequencies of mIPSC and mEPSC, which indirectly described the cellular E/I balance, was significantly increased in *Ambra1^+/gt^* females compared to control littermates, while male mice showed a similar trend without significance (Figure 5E). These data indicate that the ratio of excitatory and inhibitory inputs into single cells can be a critical factor in network homeostasis (Huang et al., 2021; Xue et al., 2014) rather than the change in the number of overall inputs themselves.

**Figure 5:**
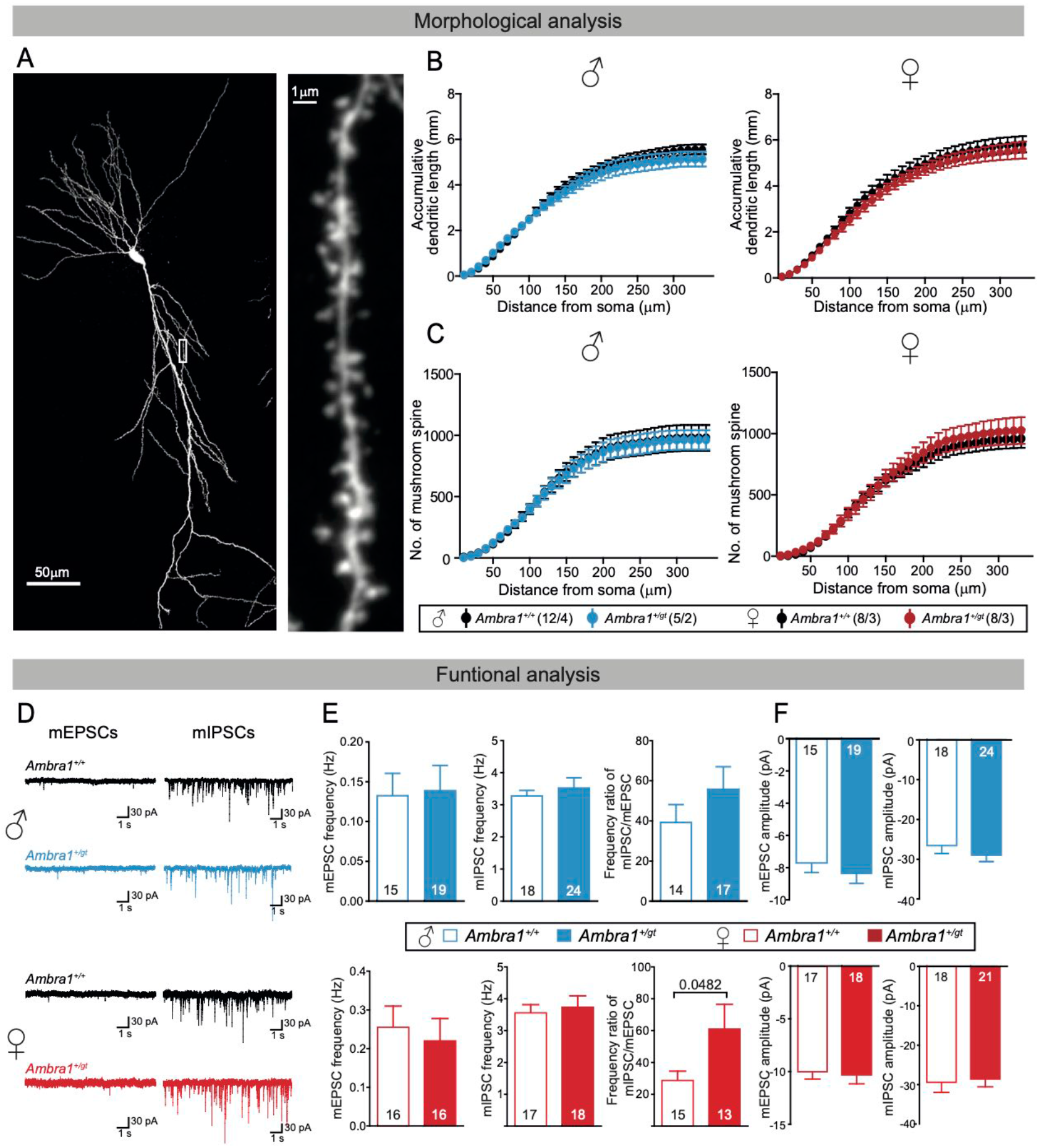
Imbalance of excitatory and inhibitory inputs upon *Ambra1* heterozygous mutation in female-specific manner. **A** Example images of CA1 pyramidal neuron. **B, C** Accumulative dendritic length (**B**) and number of mushroom-shaped spines (**C**) were plotted along the distance from soma Numbers written in brackets next to the legends represent neuron numbers/animal numbers. **D** Representative traces of mEPSC with 10μM bicuculine and mIPSC with 10μM NBQX from both genotypes. **E, F** The frequencies of mEPSC and mIPSC, frequency ratio of mIPSC/mEPSC (**E**), and amplitudes of mEPSC and mIPSC (**F**) in two genotypes of male (upper) and female (lower) mice. Numbers written within bars represent neuron numbers acquired from 4-5 animals per group. All experiments were performed at 4 weeks old. The spots and bars represent mean ± S.E.M and statistical analysis was performed by two-way ANOVA (**B, C**) and two-tailed unpaired t-test (**E, F**) with significance level p < 0.05.

Subtle phenotypes of synaptic function by *Ambra1* heterozygous mutation lead to further analysis of synaptic release by the absence of *Ambra1* gene. We accessed the synaptic release property of glutamatergic autaptic neurons cultured from cortex of *Ambra1^+/+^, Ambra1^+/gt^* and *Ambra1^gt/gt^* embryonic littermate in both sexes, at embryonic day 14.5 just before embryonic lethality (Figure 6). Since it was unlikely to obtain three genotypes of both sexes from the same littermates, the data from mutant neurons were normalized by the ones of *Ambra1^+/+^* neurons from their littermates.

**Figure 6:**
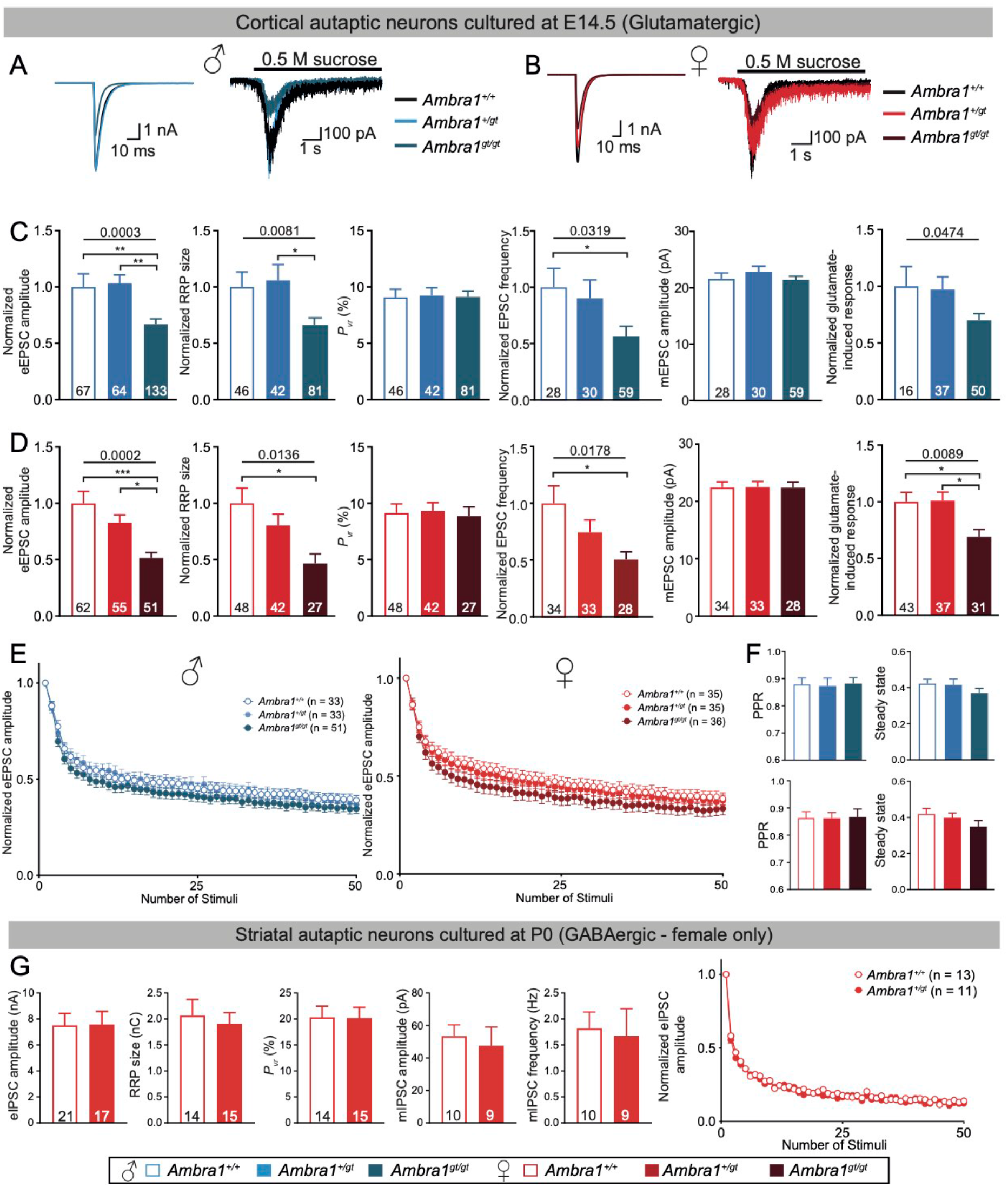
Aberrant synaptic transmission in the absence of *Ambra1* gene, regardless of sex. **A, B** Sample traces of eEPSC and 0.5 M sucrose-induced response from glutamatergic autaptic neurons from E14.5 hippocampal-like embryonic brain of *Ambra1^+/+^, Ambra1^+/gt^* and *Ambra1^gt/gt^* in males (**A**) and females (**B**). **C, D** The bar graphs of normalized eEPSC amplitude, normalized RRP size, *Pvr*, normalized mEPSC frequency, mEPSC amplitude and 100μM glutamate-induced response between three genotypes in males (**C**) and females (**D**), separately. **E** Short-term synaptic depression was monitored after application of 50 stimuli at 10 Hz. **F** Comparison of normalized eEPSC amplitude acquired from 2^nd^ stimuli (Paired pulse ratio (PPR), left) and averaged during the steady state (36-40^th^ stimuli, right) from 10 Hz stimulation experiment (**E**). **G** The bar graphs of eIPSC amplitude, RRP size, *Pvr* and amplitudes and frequency of mIPSC and a dot graph of short-term synaptic depression by 50 stimuli at 10 Hz in GABAergic autaptic neurons from P0 striatum of *Ambra1^+/+^* and *Ambra1^+/gt^* females. Neuron numbers of each group are written within the bar or next to the legends from 2-4 independent experiments and the bars and dots in graphs are presented as mean ± S.E.M. Statistical analysis of bar graphs from three groups (**C, D**, and **F**) was performed by one-way ANOVA followed by Bonferroni or Tukey post-hoc test showing significance as asterisk (*, p≤0.05; **, p≤0.01; ***, p≤0.001) and from two groups (**G**) by two-tailed unpaired t-test with significance below 0.05. Short-term synaptic depression (**E, G**) was analyzed by Repeated Measures of ANOVA.

Evoked EPSC (eEPSC) amplitude and total number of synaptic vesicles ready to release, called readily releasable pool (RRP), in *Ambra1^gt/gt^* neurons were reduced to ~67% in male and ~51% in female, respectively, of control values without change in mEPSC amplitudes and vesicular release probability (Pvr) (Figure 6A-D), suggesting that Ambra1 is essential for the activity of excitatory synapse. Interestingly, even in these cultured neurons grown independently and separately, the most prominent alteration depending on sex is that the eEPSC and RRP sizes in female heterozygous neurons, exhibited a decreasing trend, but not in males (Figure 6A-D). Synaptic plasticity upon 10 Hz stimuli was similar between three genotypes (Figure 6E-F). Interestingly, both sexes exhibited approximately 30% of reduction in response to exogeneous application of glutamate in *Ambra1^gt/gt^*, which is different from the change in synaptic responses (Figure 6C-D).

We tested whether the reduction of eEPSC amplitude is due to defects in synaptogenesis by the absence of Ambra1. Using immunofluorescent staining of vGluT1 and PSD95 to label glutamatergic pre- and postsynapses in each autaptic neuron, the number of presynaptic and postsynaptic puncta were similar between genotypes, and Mander’s overlapping coefficient between vGluT1 and PSD95 signals were comparable between genotypes, indicating that the spatial integrity of pre- and postsynapses were unaffected by Ambra1 (Figure S3). This indicates, even in cultured single neurons, sex-dimorphic changes in synaptic release were detected without alteration in synaptic number, as data in acute brain slices. In order to find out whether the phenotype also occurs in inhibitory input, the similar analysis was performed in autaptic GABAergic neurons cultured from the striatum of female mice at postnatal day 0 (Nair et al., 2013). Interestingly, the release machinery and synaptic GABA receptor cluster was not changed in *Ambra1^+/gt^* neurons (Figure 6G). Our data addressed that *Ambra1* heterozygous mutation produces a pronounced effect restricted to glutamatergic release in females without any environmental factors.

## Discussion

We characterized the consequence of *Ambra1* heterozygous mutation, known to induce female-specific ASD, by multiple level of analysis including behavioral, biochemical, morphological, and electrophysiological approaches. The remarkable and distinct finding in our study is that the reduced power of γ-oscillations and increased susceptibility to seizure in *Ambra1* heterozygous mice, two proxies of E/I imbalance, occurs exclusively in females (Figure 3). To understand the mechanism underlying this imbalance, we considered the possible contribution of autophagy that may serve as a bridge linking Ambra1 and ASD. This is because autophagy has been known to have a profound relationship with neurodegenerative diseases caused by the accumulation of harmful proteins and damaged organelles and neuronal development diseases such as ASD caused by impairment of neurodevelopmental processes including neurogenesis, neuronal differentiation and synaptic remodeling/function (Gkogkas et al., 2012; Kuijpers et al., 2021; Tang et al., 2014).

Our previous study demonstrated a marked brain enlargement in female *Ambra1^+/gt^* mutant (Mitjans et al., 2017), which is a common feature in human ASD (Courchesne et al., 2011). And Ambra1 protein is also known to be involved in proliferation/apoptosis and its absence induced brain overgrowth in mouse embryos (Maria Fimia et al., 2007). It further strengthened the autophagy hypothesis to support the ASD seen in the *Ambra1^+/gt^* and led us to speculate that altered populations of different cell types or overgrowth of neuronal morphology, such as dendritic arborization or spine density, may disturb E/I balance and contribute to brain overgrowth.

First of all, the current study displays a novel fact that Ambra1 in the brain is only present in neurons, including synapses and it limited the target of Ambra1 function (Figure 1). Moreover, the number of glutamatergic, GABAergic and PV-expressing neurons, dendrite complexity, and spine density is unaltered in *Ambra1^+/gt^* hippocampal region, indicating no critical effect on the neurogenesis by *Ambra1* heterozygous mutation (Figure 4A-G, Figure 5A-C and Figure S1-2). Crucially, no significant difference is found in autophagic activity between *Ambra1^+/gt^* and *Ambra1^+/+^* brains (Figure 4H-I), which could be supported by previous data that ~50% of autophagic activity was still observed in *Ambra1^gt/gt^* embryonic brains (Maria Fimia et al., 2007). Therefore, it can be inferred that the cause of ASD, probably E/I imbalance in *Ambra1^+/gt^*, is far from autophagy activity, which has been known to directly affects axon and synapses (Cheng et al., 2015; Soukup et al., 2016; Soykan et al., 2021; Wang et al., 2015).

The oscillatory activity in the neural network is a comprehensive signal of global E/I. To further dissect the cellular substrates underlying the alteration of this E/I signal, we focused on functional synapses rather than morphological ones. The possible cause of E/I imbalance, occurring only in female *Ambra1^+/gt^* brain slices, is a slight decrease in mEPSC frequency (Figure 5E). This moderate reduction at single cell level, through altering ratio with mIPSC frequency, can be accumulated at the neuronal network level, which contributes as the basis for inducing ASD. That is, even small changes in the number of functional synapses or synaptic activity can cause the E/I imbalance. The finding is highly reminiscent of previous study for *Nlgn4* knockout mice, as a construct-valid and face-valid mouse model of ASD, proposing that the accumulation of subtle local changes in synaptic function yields pronounced perturbation in global network activity (Hammer et al., 2015). We projected the “little things make great things” into our data. However, it was still not sufficient to explain the mechanism by which ASD occurring in *Ambra1* heterozygous mutation appear only in female.

The female-specific behavioral phenotypes had to consider the influence of specific environments *in vivo*, such as unique hormones. To study the neuronal intrinsic change by genetic factors limited to females, eliminate the external effects coming from *in vivo* conditions, and deconvolute the diluted results from E/I input measurement, we specifically analyzed the cultured neurons from very early embryonic day 14.5 just before *Ambra1^gt/gt^* embryos death (Maria Fimia et al., 2007). As the intrinsic activities of cultured *Ambra1^+/+^, Ambra1^+/gt^*, and *Ambra1^gt/gt^* neurons could be compared simultaneously, it enables us to understand the substantial role of Ambra1 in neurons. As a result, regardless of sex, Ambra1 deficiency had no effect on neuronal development and synapse formation, and decrease selectively the number of functional glutamatergic synapses. Moreover, surprisingly, the cultured *Ambra1^+/gt^* neurons also showed a female-specific decrease in eEPSC size, as the change in mEPSC frequency in *Ambra1^+/gt^* female brain acute slice (Figure 6A-D). Thus, ASD caused by E/I imbalance in *Ambra1* heterozygous mutation is a neuron-intrinsic property of Ambra1 by sex difference without any environmental factors. And our previous study, the decrease in relative Ambra1 expression level in female *Ambra1^+/gt^* brain, compared to one in male (Dere et al., 2014), may help to understand the female-specific synaptic phenotype. Thus, we can suggest that the size of EPSCs would be determined according to the expression level of Ambra1 protein.

In *Ambra1 ^gt/gt^* neurons, the comparable mEPSC amplitudes and the eEPSC or mEPSC frequencies reduced in half lead to the novel fact that there are Ambra1-dependent and - independent synapses. In particular, it can be speculated that Ambra1 deficiency makes Ambra1-dependent synapses into silence (Figure 6). And, although Ambra1-dependent synapses maintain silence, the fact that the reduction of glutamate-induced response is less than that of synaptic responses such as eEPSC, mEPSC frequency and RRP size raise two possibilities (Figure 6). Firstly, depletion of functional synaptic receptors that may be induced by Ambra1 deficiency can lead to an increase in the number of extrasynaptic receptors. The other possibility is that the *Ambra1^gt/gt^* neurons display the complete ablation of synaptic release in Ambra1-dependent synapses. To elucidate the impairment of synaptic release and its relationship to synaptic receptors, we recall our previous work (Sigler et al., 2017). The number of functional synaptic glutamate receptors was reduced by approximately 40% in Munc13-deficient synapses in which synaptic transmission from presynaptic terminal is completely impaired. So, the difference between reduction ratios in synaptic parameters such as the eEPSC size and mEPSC frequency, and glutamate induced responses in *Ambra1^gt/gt^* can be attributed to the complete impairment of glutamate release. In order to support the two hypotheses, it can be inferred that the Ambra1-dependent synapses accounts for about 50% of the total synapses. Considering our data evaluating the expression levels of synaptic receptors and scaffolding proteins (Figure S1), we highly appreciate the latter possibility. We figured out that the most critical factor of E/I imbalance in *Ambra1* heterozygous brain is the contribution of Ambra1 in the activity of glutamatergic synapses, regardless of autophagy. In the end, we discovered that the novel function of Ambra1 in neuronal cells play an important factor in ASD manifestation.

We conclude that Ambra1, localized specifically in neuronal cells, intrinsically triggers excitatory synaptic activity, and that its sex-dimorphic expression of protein level modulates the degree of glutamate release depending on sex (Dere et al., 2014). However, the mechanism for the sex-dimorphic expression of Ambra1 protein level is still an important piece to be studied further. This sex-dimorphic reduction of synaptic transmission by *Ambra1* haploinsufficiency might manifest female-specific phenotypes, such as autistic-like behaviors, increased seizure propensity and aberrant gamma oscillations (Dere et al., 2014), which provides us important insight on the neural substrate of E/I balance related with ASD and epilepsy.

## Materials and methods

### Animals

*Ambra1* mutant mice were described previously(Maria Fimia et al., 2007). Wild-type (WT, *Ambra1^+/+^*) and heterozygous *Ambra1^+/gt^* (Het) littermates of both sexes with a >99% C57BL/6N genetic background were obtained by interbreeding male *Ambra1^+/gt^* and female WT C57BL/6N mice and used for all experiments on postnatal animals. For neuronal cultures, *Ambra1^+/+^, Ambra1^+/gt^*, and *Ambra1^gt/gt^* (KO) littermate embryos were obtained by interbreeding male and female *Ambra1^+/gt^* mice. All experiments were carried out in agreement with the guidelines for the welfare of experimental animals issued by the Federal Government of Germany and Max Planck Society.

### Genotyping

Genomic DNA for genotyping was extracted from tail tips of 2-3 week-old offspring or embryos using NucleoSpin Tissue kit (Machery-Nagel GmbH & Co. KG). WT and KO *Ambra1* alleles and Y chromosomes of offspring were detected by polymerase chain reaction (PCR) of genomic DNA. PCR analyses of *Ambra1* KO gene were performed as described previously(Dere et al., 2014) using GoTaq^®^ G2 Flexi DNA polymerase (Promega). For *Ambra1* WT allele, GoTaq^®^ G2 Flexi DNA polymerase (Promega) with forward primer, 5’-AAC TGA ACC TGG GTT CTT TGA A-3’ and reverse primer 5’-GAA AAG CTC CCC ATC TTT TCT T-3’ were used to generate a 0.5 kb fragment (95°C/5 min, 35 cycles with 95°C/30 s, 57°C/45 s, 72°C/105 s, and 72°C/ 10 min). For sex determination of embryos, PCR analyses of Y chromosomes were performed using GoTaq^®^ G2 Flexi DNA polymerase, forward primer 5’-GGT GTG GTC CCG TGG TGA GAG-3’, and reverse primer 5’-GAG GCA ACT GCA GGC TGT AAA ATG-3’ to generate a 270 bp fragment (94°C/1 min, 33 cycles with 94°C/1 min, 63°C/30 s, 72°C/30 s, and 72°C/7 min). PCR products were analyzed on a 1.5% agarose gel in Tris-Acetate-EDTA buffer, which were stained with HDGreen^®^ Plus Safe DNA Dye (Intas).

### mRNA expression of Ambra1 from data of Allen Brain Atlas

Data of mRNA expression level in different brain regions were extracted from Allen Mouse Brain Atlas (Figure 1a, http://mouse.brain-map.org/) (Ju et al., 2020; Lein et al., 2007). mRNA expression level in regions of interests (ROIs) of in situ hybridization was calculated by multiplying expression density and intensity.

### Histological and Immunohistochemical Analyses

Mice were perfused transcardially with Ringer solution followed by 4% paraformaldehyde (PFA) in 0.1 M phosphate buffer (PBS, pH=7.4). Brains were post-fixed at 4°C in 4% PFA in PBS for 2 h for X-galactosidase (X-gal) histochemical staining, or post-fixed overnight, followed by cryo-protection in 30% sucrose solution in PBS and in liquid nitrogen for immunohistochemistry.

X-gal histochemical staining was performed with brains of 9 weeks old mice. Coronal brain sections (50 μm) were cut using Leica VT1000S vibrotome (Leica Biosystems) and incubated overnight in the dark at 37°C in X-gal solution containing 5mM K3[Fe(CN)6]. 5mM K4[Fe(CN)6], 2mM MgCl_2_ and 1.2 mg/mL 5-bromo-2-chloro-3 indoyl-b-D-galactopyranoside (X-gal) in PBS, rinsed three times in PBS, and mounted with Aqua-Poly/mount (Polyscience). Digital images were obtained using an Axiophot microscope (Carls Zeiss Microscopy GmbH).

*Ambra1* WT and Het female mouse brain were cut into coronal sections (30 μm) with Leica CM1950 instrument (Leica Biosystems). Sections were blocked with 10% normal horse serum (NHS) and 0.2% Triton-X-100 in PBS for 1 h at room temperature (RT). PBS with 5% NHS and 0.2% Triton-X-100 was also used for the primary and secondary antibody dilution. Incubation of the primary antibodies was carried out at 4°C for 1-3 nights, followed by incubation of secondary antibodies (1:500) for 2 h and DAPI (1:10,000, D9542, Sigma-Aldrich) in PBS for 5 min at RT. Washing was performed between every step and sections were mounted using Aqua-Poly/mount. The following antibodies were used for immuohistochemistry: mouse anti-β-gal (Z3781, Promega), chicken anti-NeuN (266006, Synaptic Systems), rabbit anti-IBA1 (019-19741, Wako), rabbit anti-Olig2 (AB9610, Chemicon), rabbit anti-GFAP (G5601, Promega), guinea pig anti-Ctip 2 (325005, Synaptic Systems), mouse anti-GAD67 (MAB5406, Chemicon), rabbit anti-PV (PV27, Swant), Alexa-Fluor 488 donkey anti-mouse IgG (A21202, Invitrogen), Alexa-Fluor 488 donkey anti-chicken IgG (703-546-155, Jackson ImmunoResearch), Alexa-Fluor 488 goat anti-chicken IgG (A-11039, Thermo Fisher Scientific), Alexa-Fluor 555 goat anti-guinea pig IgG (A-21435, Thermo Fisher Scientific), Alexa-Fluor 555 goat anti-mouse IgG (A-21424, Thermo Fisher Scientific), Alexa-Fluor 594 goat anti-mouse IgG (115-585-003, Jackson ImmunoResearch), Alexa-Fluor 594 donkey anti-rabbit IgG (A-21207, Invitrogen) and Alexa-Fluor 633 goat anti-rabbit IgG (A-21071, Thermo Fisher Scientific). Leica TCS SP5 confocal microscope (Leica Biosystems) was used to scan anatomically matched sections using 0.5 μm z-step and a 20x objective lens. For counting cell numbers, the dorsal part of hippocampus (Bregma −1.34 to −2.54 mm posterior) was used bilaterally in each animal (12-14 hippocampi per 1-2 animal). Image stacks were further processed by Image J and quantification of CTIP2+, GAD67+ and PV+ were done using Imaris 7.5.1 and manually. Cell density was obtained by dividing the number of each cell type by the total volume of hippocampal region in mm^3^.

### Protein Extraction and Measurement

Cortices of 6-weeks old mice and hippocampi of 4-weeks old mice were dissected in cold 0.32 M sucrose solution with protease inhibitors (0.1 μM Aprotinin, 50 μM Leupeptin, 0.2 mM PMSF) and homogenized using glass-teflon homogenizer (900 rpm, 10 strokes). Cortical homognates of male WT and female WT and Het mice at 6 weeks of age were used for purification of synaptic membrane proteins (Fig. 1f). All centrifugations were performed with Beckman TL-100 Ultracentrifuge (Beckman Coulter) at 4°C. Cortical homogenates were layered on a discontinuous gradient of 0.85 M, 1.0 M, and 1.2 M sucrose solutions. After centrifugation at 82,500 g for 2 h, the supernatant above 0.85 M sucrose layer and the pellet were kept as soluble fraction (S) and mitochondria-enriched fraction (P2D), respectively. The interface fractions between 0.32 M and 0.85 M sucrose, between 0.85 M and 1.0 M sucrose, and between 1.0 M and1.2 M were collected as myelin-enriched fraction (P2A), ER-Golgi-enriched Fraction (P2B), and synaptosome fraction (P2C), respectively. The P2C fraction was diluted with 0.32 M sucrose solution with protease inhibitors and centrifuged at 100,000 g for 20 min. After centrifugation, the resulting pellet was resuspended in 2.5 mL of 6 mM Tris-Cl, pH 8 and incubated on ice for 45 min for osmotic shock. After centrifugation at 32,800 g for 20 min, the supernatants were collected as synaptic cytoplasm and crude synaptic vesicle (SC/CSV) fractions, and the pellets were resuspended as crude synaptic membrane (CSM) fraction with 3 mL of 0.32 M sucrose solution with protease inhibitors. CSM fractions were applied to a discontinuous gradient of 0.85 M, 1.0 M, and 1.2 M sucrose solutions and centrifuged at 82,500 g for 2 h. The interface fraction between 1.0 M and 1.2 M sucrose was harvested as pure synaptic membrane fraction (SM) and diluted in 0.32 M Sucrose solution with protease inhibitors, followed by centrifugation at 100,000 g for 20 min. The resulting pellet was resuspended in 500 μL of 6 mM Tris-Cl pH 8.0. Purified fractions were stored at −80°C. Protein concentrations in various samples were measured using Bradford method (Bio-Rad) according to the manufacturer’s instructions.

### Western Blotting

SDS-PAGE and protein transfer to nitrocellulose membranes were performed according to standard procedures.(Laemmli, 1970; Towbin et al., 1979) After transfer, the membranes were washed with ultra-pure water and incubated with Memcode Reversible Protein Stain Kit (Thermo Fisher Scientific) according to the manufacturers’ protocol. Membranes were washed and incubated in blocking buffer (5% milk powder in Tris-based saline with 0.05 % Tween-20, TBST) for 1 h at RT. followed by incubation with primary antibodies diluted at 1:1000 in blocking buffer for 3 h at RT. Membranes were then incubated with primary antibodies (1:1000) and secondary antibodies (1:5000) with washing three times with TBST for 15 min each between. Protein signals were detected with Odyssey Infrared Imaging System (LI-COR Biosciences GmbH) and quantified using the Image-Studio Software (Odyssey System; LI-COR Biosciences GmbH) with normalization to total protein assessed by Memcode staining. The following antibodies were used for Western blotting: mouse anti-PSD95 (ab2723, Abcam), mouse anti-Gephyrin (147111, Synaptic Systems), rabbit anti-GluR1 (PC246, Calbiochem), rabbit anti-GluR2 (182103, Synaptic Systems), mouse anti-NMDAR1 (114011, Synaptic Systems), rabbit anti-GuR6/7 (04-921, Mllipore), rabbit anti-GABAARα1 (Ab5592, Chemicon GmbH), rabbit anti-GABAARγ2 (Ab82970, Abcam), rabbit anti-vGluT1 (135302, Synaptic Systems), mouse anti-ß-Tubulin (T4026, Sigma-Aldrich), IRDye680-anti-mouse IgG (926-68070, LI-COR Biosciences GmbH), IRDye800-anti-rabbit IgG (926-32211, LI-COR Biosciences GmbH).

### Pentylenetetrazol (PTZ)-Induced Seizures

Seizure activity was induced in awake 12-13 weeks-old Ambra1 WT and Het mice of both sexes by a single intraperitoneal (i.p.) injection of 50 mg of Pentylenetetrazol (PTZ; P6500, Sigma-Aldrich) per 1 kg of body weight. After injection, mice were observed closely for 30 min in a small clear home cage. Four phases of behavioral response to PTZ injection were defined as follows: (1) Hypoactivity; decrease in mobility until the animal arrests in a crouched posture. (2) Partial clonus (PC), clonic seizure activity in face, head, and forelimbs. (3) Generalized clonus (GC); sudden loss of upright posture, whole body clonus including all four limbs and tail, rearing and autonomic signs. (4) Tonic-clonic (TC) (maximal) seizure; generalized seizure with tonic hind-limb extension followed by death. The latency to GC and the seizure score, which is calculated from the latencies to PC, GC, and TC in seconds by equation [Seizure score = 1000 / (0.2 * PC latency + 0.3 * GC latency + 0.5 * TC latency)] were used as measures(Wojcik et al., 2013).

### Electrophysiological recordings

Four weeks old WT and *Ambra1* Het mice were anesthetized with Isofluorane and decapitated. During the entire procedure, carbogen gas (95% oxygen and 5% carbon dioxide) keeps applied in solutions. The whole brain was immediately transferred to cold slicing solution (230 mM Sucrose, 26 mM NaHCO_3_, 1 m KH_2_PO_4_, 2 mM KCl, 2 mM MgCl_2_x6H_2_O, 10 mM Glucose, 0.5 mM CaCl_2_). To get hippocampal slices for evoke field excitatory postsynaptic potentials (fEPSP) recording, hippocampi were isolated carefully and cut transversally at 300 μm thickness using a McILWAIN tissue chopper (Molecular Devices, LLC). For acute brain slices, sagittal sections at 5° angle tilted to the midline with 300 μm thickness were obtained inside the same slicing solution at 4°C using Leica VT1200S vibrotome (Leica Biosystems). Slices were immediately transferred to a chamber filled with artificial cerebrospinal fluid (ACSF; 120 mM NaCl, 26 mM NaHCO_3_, 1 m KH_2_PO_4_, 2 mM KCl, 2 mM MgCl_2_x6H_2_O, 10 mM Glucose, 2 mM CaCl_2_) at 37°C for 1h 40min for hippocampal slices and at 37°C for 20 min for acute brain slices. After recovery, slices were kept at RT.

For fEPSP measurement, the hippocampal slices were placed in interface recording chamber (Harvard Apparatus) with continuous flow of carbogen-supplied ACSF at 30°C. An electric stimulation was applied with 100 μs duration time by concentric metal bipolar electrode (FHC) on the *Stratum radiatum* of Schaffer collaterals. Recording electrode (2-3 MΩ) was pulled from thin-walled borosilicate glass capillaries, filled with ACSF, and positioned on the *Stratum radiatum* of CA1 area.

For kainite-induced gamma oscillation recording, acute brain slices were placed on interface recording chamber with application of ACSF at 33°C. Recording electrode (2-3 MΩ), filled with ACSF, was placed in the CA3 pyramidal layer of hippocampus. For each slice, baseline field potentials were recorded for 30 min in ACSF, followed by recording of oscillatory field potentials in gamma-range induced by 100 nM kainic acid (BN0281, BIOTREND Chemikalien GmbH) in ACSF for 30 min. After this recording phase, the electrode was slightly re-positioned to acquire the maximum power of gamma oscillation for another 10 min. Recordings were acquired by Multiclamp 700B amplifier and Digidata 1440A (Molecular Devices, LLC.) and data were analyzed using AxographX (Axograph), as previously described(Ripamonti et al., 2017).

For whole-cell recordings, the somata of hippocampal CA1 pyramidal neurons in acute brain slice were whole-cell voltage clamped at −70 mV by recording electrode (2.5-3.5 MΩ) containing internal solution with 100 mM KCl, 50 mM K-gluconate, 10 mM HEPES, 4 mM ATP-Mg, 0.3 mM GTP-Na, 0.1 mM EGTA, 0.4% biocytin, pH 7.4, 300 mOsm. The external solution was carbogen-saturated ACSF. Miniature excitatory and inhibitory post-synaptic currents (mEPSCs/mIPSCs) were recorded in the presence of 1 μM TTX, mixed with 10 μM bicuculine methiodide for measuring mEPSCs or with 10 μM NBQX for measuring mIPSCs with washing with 1 μM TTX for 15 mins between. An EPC-10 amplifier with Patchmaster v2X80 software was used for data acquisition (HEKA/Harvard Bioscience). Subsequently, slices were fixed using 4% PFA in PBS for two hours at RT and washed by PBS.

### Immunohistochemistry of Biocytin-filled Neurons

After being washed in PBS, blocked and permeabilized for 1h with blocking solution (5% NGS and 0.5% Triton X-100 in PBS), fixed brain slices obtained from patching were stained with Alexa-Fluor-555-labeled streptavidin (1:1000; S32355, Thermo Fisher Scientific.) and DAPI (1:10,000) in blocking solution. After washing, the slices were mounted on glass slides and covered with cover slips in Aqua-Poly/Mount. CA1 pyramidal neurons in hippocampus were scanned using a Leica SP5 confocal microscope with 100 x/1.44 NA oil objective at 0.126 μm z-intervals. The basal and apical part of pyramidal neurons 3D Gaussian-filtered (σ_x,y_ 0.7, σ_z_ 0.7,), using custom-written macros to handle the large data sets. Only cells with a pyramidal shape and a location in CA1 were used for further analysis.

### In Utero Electroporation and Immunohistochemistry for Sholl Analysis

E14.5 mouse embryos from WT mothers bred with Het males were subjected to IUE (permit number 33.19-42502-04-13/1052), as previously described(dal Maschio et al., 2012; Hsia et al., 2014). DNA solution with pFUGW (0.1 mg/mL) and pCX::myrVENUS (0.1 mg/mL) for the myrVenus construct(Lois et al., 2002; Rhee et al., 2006) were used to sparsely label CA1 pyramidal neurons in hippocampus.

*In utero* electroporated mice were perfused, and brains were post-fixed and cryo-protected (15%-30% sucrose in PBS) at P28. Coronal brain sections (230 μm thickness) at −1.06 mm to −2.46 mm from Bregma were collected using a Leica VT 1000S vibrotome. For immunofluorescence staining, PFA was quenched by 1 mg/mL NaBH_4_ in PBS for 5 min followed byh thorough washing in PBS. Brain sections were incubated in blocking solution (5% normal goat serum (NGS) and 0.5 % Triton-X-100 in PBS) for 1 h at RT followed by incubation in 0.2% Tween-20 and 10 μg/mL heparin in PBS for 1.5 h to improve the penetration of antibody in thick brain sections. The blocking solution was used for diluting primary and secondary antibodies (1:1000 dilution). The sections were incubated with polyclonal rabbit anti-GFP antibody (598, MBL) for 4 days at 4°C and with Alexa-Fluor (AF) 488 goat anti-rabbit IgG (R37116, Thermo Fisher Scientific) overnight at RT followed by DAPI staining (1:10,000) and mounted on slides. Images of CA1 pyramidal neurons in hippocampus were acquired with 1.02 μm z-steps by Leica SP2 confocal microscope (Leica Biosystems) and oil-immersion 20x objective.

### Analysis of Neuron Morphology using NeuronStudio

For segmentation of entire dendritic trees and subsequent mushroom spine analysis, we used NeuronStudio (CNIC, Mount Sinai School of Medicine, New York, NY, USA)(Rodriguez et al., 2008). After reconstructing the dendritic trees, 3D Sholl analysis from this program was performed to acquire the accumulative dendritic length in every 10 μm step from the center of soma.(Sholl, 1953) Mushroom spines were automatically detected along the reconstructed dendritic tress using the NeuronStudio segmentation algorithm by keeping the suggested parameters(Rodriguez et al., 2008; Sigler et al., 2017) (Head diameter of mushroom spines: 0.35 μm) and the numbers of mushroom spines were counted every 10 μm from the center of soma.

### Autaptic Neuron Culture and Electrophysiology

Autaptic cultures of cortical glutamatergic neurons from hippocampi from E14.5 embronic brains or striatum of P0 postnatal brains were prepared according to a previously published protocol(Jockusch et al., 2007) with slight modifications. 3,000 to 3,500 cells were plated on 35 mm^2^ coverslips with astrocyte islands in Neurobasal-A Medium with supplements.

Autaptic neurons were analyzed electrophysiologically as described previously(Kawabe et al., 2010) at day in vitro (DIV) 10-16. Autaptic neruons were whole-cell voltage clamped at −70 mV with a MultiClamp700B amplifier (Axon Instruments, Molecular Devices) under the control of the Clampex program 10.1 (Molecular Devices). The internal solution for recording autaptic neurons consisted of 136 mM KCl, 17.8 mM HEPES, 1 mM EGTA, 4.6 mM MgCl_2_, 4 mM NaATP, 0.3 mM Na_2_GTP, 15 mM creatine phosphate, and 5 units/mL phospho-creatine kinase (315-320 mosmol/L), pH7.4. Extracellular solution contained 140 mM NaCl, 2.4 mM KCl, 10 mM HEPES, 10mM glucose, 4 mM CaCl_2_, and 4 mM MgCl_2_ (320 mosmol/L), pH 7.3. All chemicals were purchased from Sigma-Aldrich (Sigma-Aldrich) unless mentioned otherwise.

Evoked post synaptic currents (PSCs) were measured by depolarization of neurons from −70 to 0 mV for 2 ms. Readily releasable vesicle pool size (RRP) was recorded by measuring PSC in response to application of 0.5 M hypertonic sucrose in extracellular solution. *P_vr_* was calculated by dividing the charge transfer during an action potential-evoked response by the charge transfer measured during a response to hypertonic solution. Additionally, EPSC amplitudes were measured after application of 50 stimuli at 10 Hz to assess short-term depression. Glutamate-induced response was measured by focal application of 100 μM glutamic acid (Sigma) in order to study the cell surface expression of glutamate receptor. mEPSCs were recorded in the presence of 300 mM TTX (Tocris). Given that three genotypes of identical sex are difficult to obtain in the same litters, the data were normalized to WT mean values obtained in several sets of experiments.

### Immunofluorescent staining and analysis of pre- and post-synaptic puncta in autaptic neurons

At DIV 18-23 from two independent cell preparation, autaptic cultured neurons were fixed in a solution containing 4% PFA/4% sucrose in PBS, pH7.4, for 20 min. After washing with PBS for 3 times, cells were incubated in blocking solution containing 0.3% Triton-X-100, 10% NGS and 0.1% fish skin gelatin (Sigma) in PBS for 20 min. Neurons were incubated with primary antibodies against vGluT1 (1:1000, rabbit polyclonal, 135303, Synaptic Systems), PSD95 (1:200, mouse monoclonal, ab2723, Abcam), and MAP2 (1:500, chicken polyclonal, NB300-213, Novus biologicals) diluted in blocking solution for overnight at 4°C. After repetitive washing with PBS, neurons were incubated with secondary antibodies diluted in blocking solution, including Alexa-Fluor 488 goat anti-chicken IgG (1:1000, A-21441, Invitrogen), Alexa-Fluor 555 goat anti-rabbit IgG (1:1000, A-32732, Invitrogen) and Alexa-Fluor 633 goat antimouse IgG (1:1000, A-21052, Invitrogen) for 2 hrs at RT, followed by another rounds of washing. After DAPI staining, coverslips were mounted on slides.

Single autaptic neurons were imaged by Leica SP2 confocal microscope using 40x objective (resolution: 1024 x 1024 pixels) with 1 μm z-step and analyzed using Image J software, as referenced from previous publication(Ripamonti et al., 2017). Briefly, for the quantification of pre- and post-syanptic puncta, the fluorescent signals of vGlut and PSD95 were thresholded and binarized followed by being watersheded. The number of their puncta was analyzed using ‘Analyze particle’ option. For the colocalization of pre- and post-synaptic marker, the Manders’ overlap coefficient was calculated by Intensity Correlation Analysis plugin.

### Statistical Analysis

All data were analyzed separately for males and females. Statistical methods are described in figure legends. All statistics were performed with Excel (Microsoft), GraphPad Prism 5 software (GraphPad software) and SPSS 17 (SPSS Inc.). Data are presented as mean±S.E.M., and p-values <0.05 were considered as indicating a significant difference.

## Acknowledgment

We thank F. Benseler, for valuable advice and excellent technical support. We are grateful to the staffs at the animal facility of the Max Planck Institute for Multidisciplinary Sciences for mouse husbandry.

## Additional information

### Funding

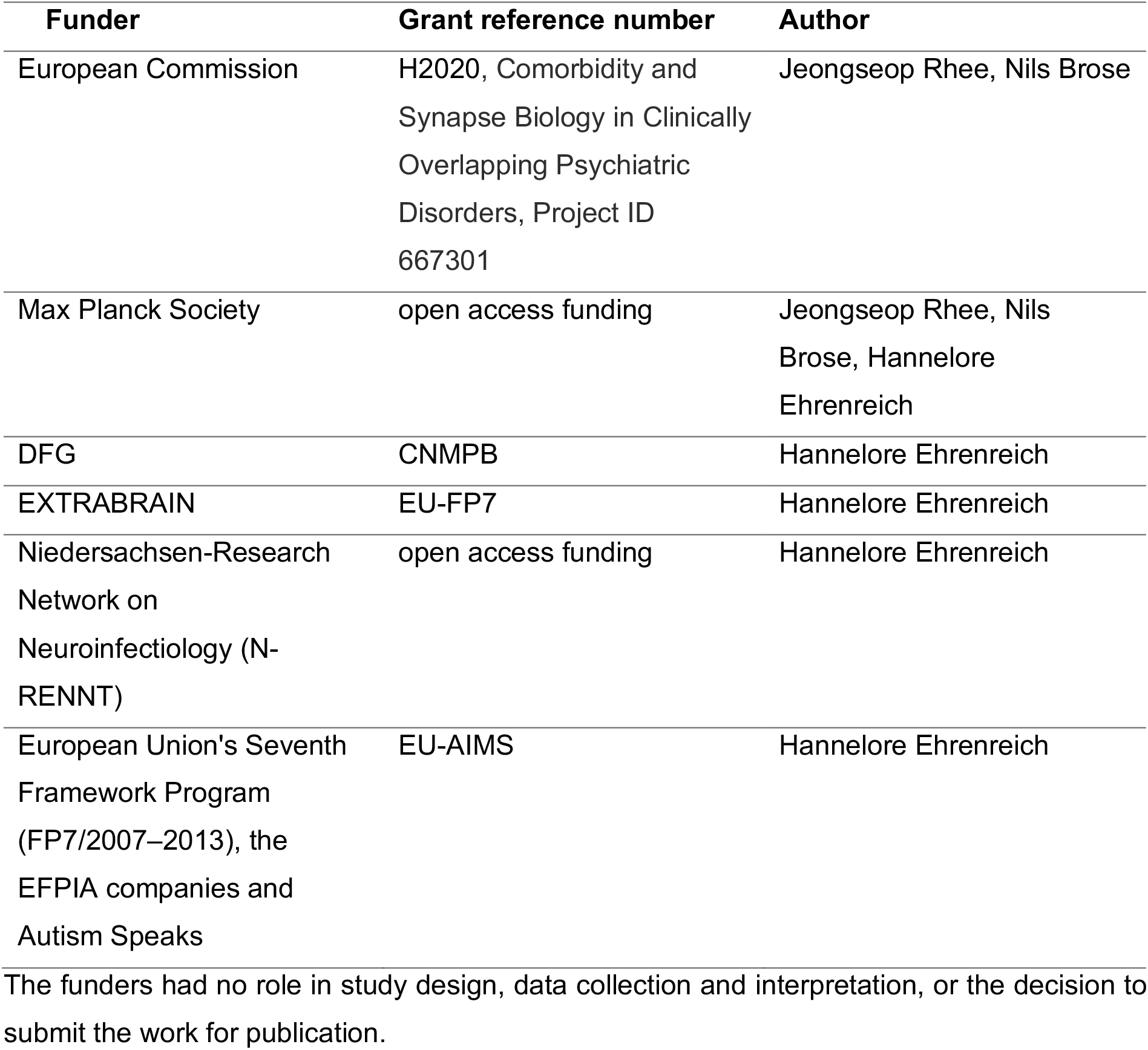

### Author contributions

Conceptualization: JSR and AJ; Methodology: JSR, AJ, BA, HJR, AS, HE, IH, MS, HK; Investigation: JSR, AJ, BA, HJR, HE, IH, MS, HK; Writing (Original draft): JS and AJ; Writing (review & editing): JS, AJ and NB; Funding acquisition: JSR, NB and HE; Resources: JSR, NB and HE; Supervision: JSR

Nils Brose (NB) is under competing interests for being a reviewing editor of this journal. Other authors declare that there is no competing of interest.

### Ethics

All experiments were carried out in agreement with the guidelines for the welfare of experimental animals issued by the Federal Government of Germany and Max Planck Society.

### Data availability

Most of data generated or analyzed during this study are included in the manuscript and supporting files.

## Supplemental Information

### Developmental features in this mouse line

A previous publication has reported alterations in synaptic plasticity, pyramidal neuron spine density and number of parvalbumin-expressing neurons in the hippocampus of adult mice (8 weeks old) restricted to females, whereas we observed no difference in our 4 week-old mice (Nobili et al., 2018). Therefore, we hypothesized that the developmental stage is a pivotal factor to explain the neural substrate underlying and the prepubertal stage is very critical time window in this mouse line. We previously reported very interesting developmental features in seizure propensity, showing opposite transition from protective response to seizure induction in female mutants of 3 weeks old to reduced survival in 13 months old (Mitjans et al., 2017). Here, we added seizure susceptibility at 12 weeks old where this transition already happened, indicating developmental progression of seizure propensity in this mouse line (Mitjans et al., 2017).

---

**Supplementary Figure S1:**
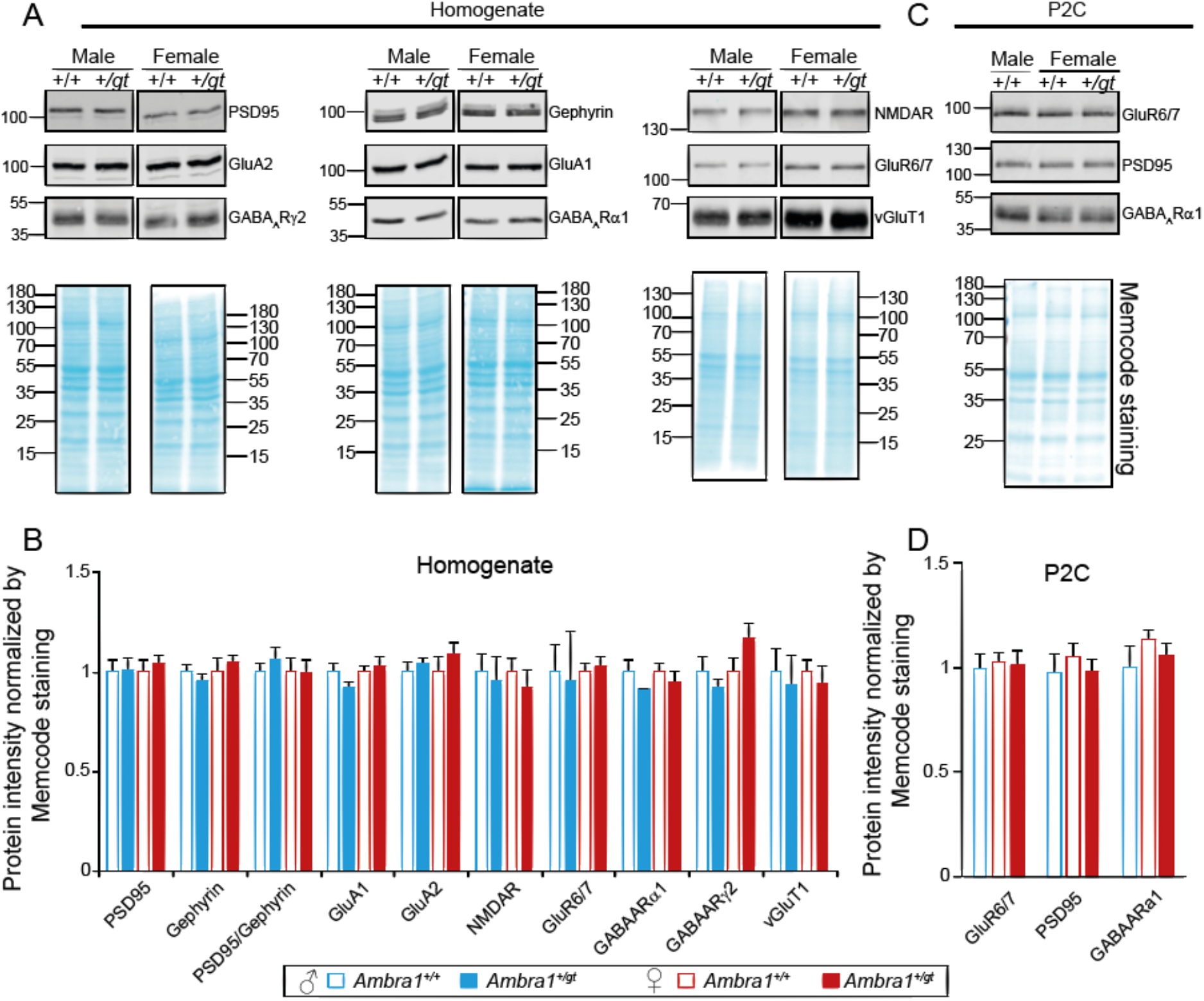
Quantification of synaptic proteins in homogenate and synaptosomes. Sample pictures of synaptic proteins (**A, C**) and corresponding Memcode staining and their quantification (**B, D**) were obtained from hippocampal homogenates of *Ambra1^+/+^* (+/+) and *Ambra1^+/gt^ (+/gt*) in 4 week-old males and females (**A, B**) and cortical P2C of 6 week-old male +/+ and female +/+ and *+/gt* (**C, D**). The intensity of synaptic proteins was normalized by corresponding MemCode staining as the amount of total proteins and normalized by the average value of male wild-type group. 3-7 mice were used per each group. The bar graphs represent mean ± S.E.M and statistical analysis between two genotypes were carried out by two-tailed unpaired t-test with significance level p < 0.05.

**Supplementary Figure S2:**
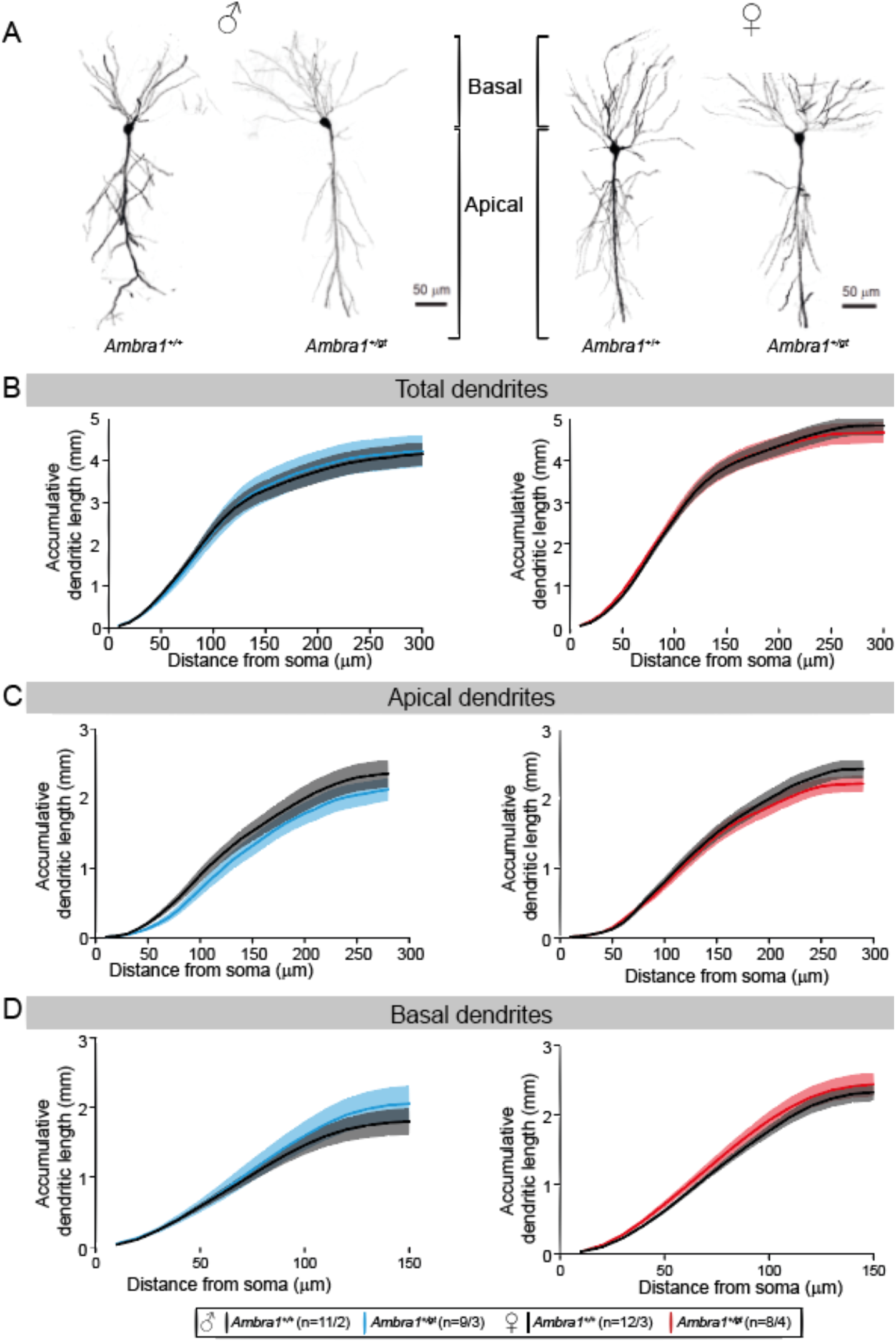
Sholl analysis from *in utero* electroporated neurons. Data from male and female mice are placed on the left and right side, respectively. A Representative picture of CA1 pyramidal neurons in hippocampus of *Ambra1^+/+^* and *Ambra1^+/gt^* mice in males and females at P28. B-D Comparison of dendritic length in whole (B), apical (C) and basal (D) parts of pyramidal neurons in every 10 μm between genotypes in male and female, separately. Neuron number/animal number are written next to the legends. Mean ± S.E.M. are presented in line and area and statistical difference was defined by p-value between genotypes from Repeated-Measures of ANOVA.

**Supplementary Figure S3:**
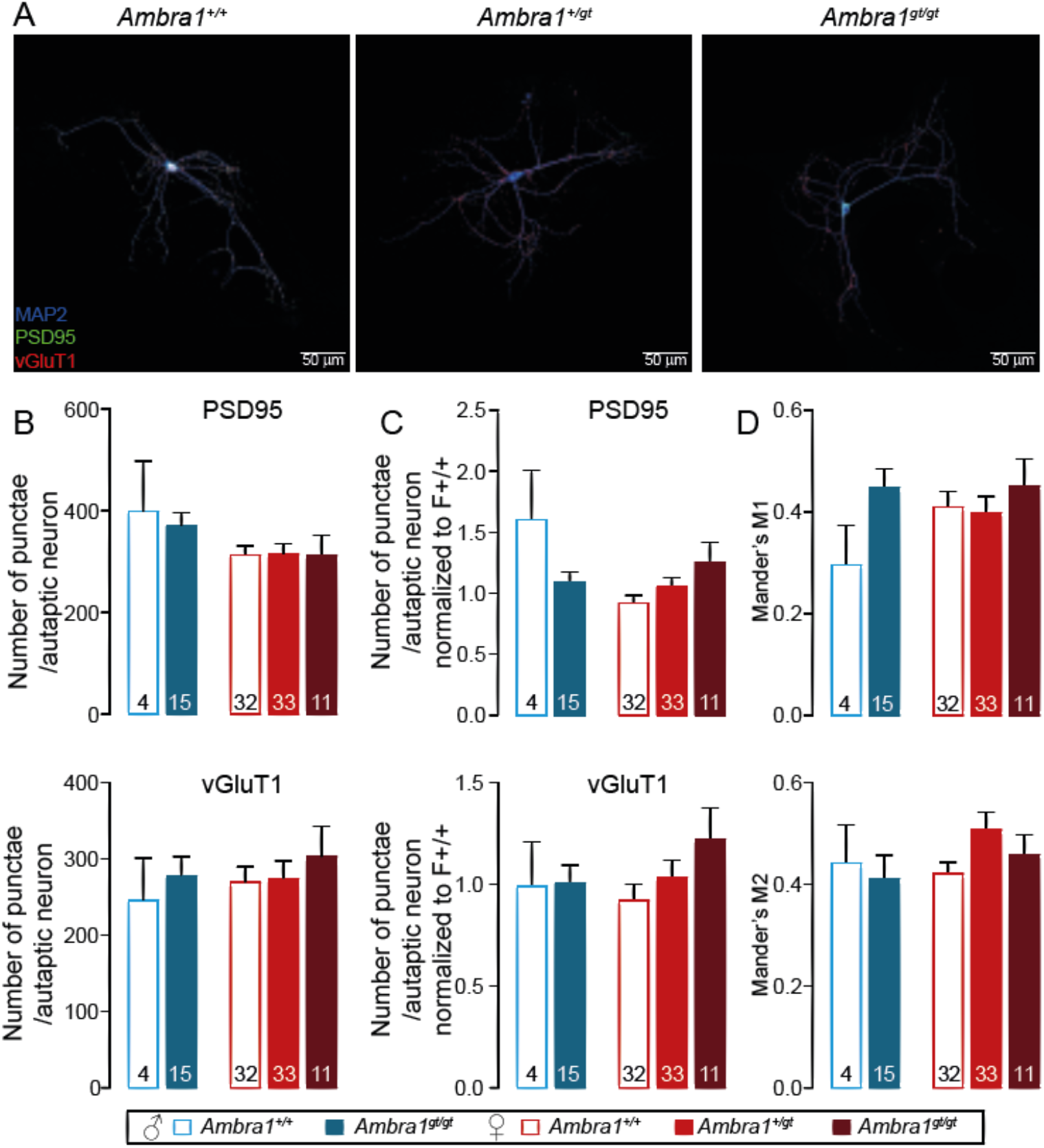
Quantification of synapse number in glutamatergic autaptic hippocampal neurons. **A** Representative images of glutamatergic autaptic neurons from *Ambra1^+/+^, Ambra1^+/gt^* and *Ambra1^gt/gt^* of E14.5 female hippocampi-like structures stained with antibodies against vGluT1, PSD95 and MAP2 at DIV18-23. **B, C** The absolute (**B**) or normalized number (**C**) of pre- (vGluT1-positive, above) or post-synaptic (PSD95-positive, below) puncta are shown. The normalization was done by dividing the average value of female *Ambra1^+/+^* cultured on the same day. **d** M1 (above) and M2 (below) coefficient of Mander’s overlapping analysis between vGluT1 and PSD95 signals. The bar graphs represent mean ± S.E.M and the numbers of analyzed neurons are written inside the bottom of each bar. Statistical analysis between genotypes were carried out by two-way ANOVA followed by post-hoc Bonferroni test with significance level p < 0.05.

## Notes

### Summary of Updates

This new version of the article contains a corrected version of panel A in Supplementary Figure S1. In the original version, the left and middle columns of this panel were inadvertently assembled using images of Memcode-stained blots from an iteration of the experiment that did not correspond to the respective Western blot images. In the corrected version of the figure, Memcode and Western blot images in each column are from the same experiments

