## Supplementary figures and images for "Regulation of excitatory presynaptic activity by Ambra1 protein determines neuronal networks in sex-dimorphic manner"

### supplementary Figure 1

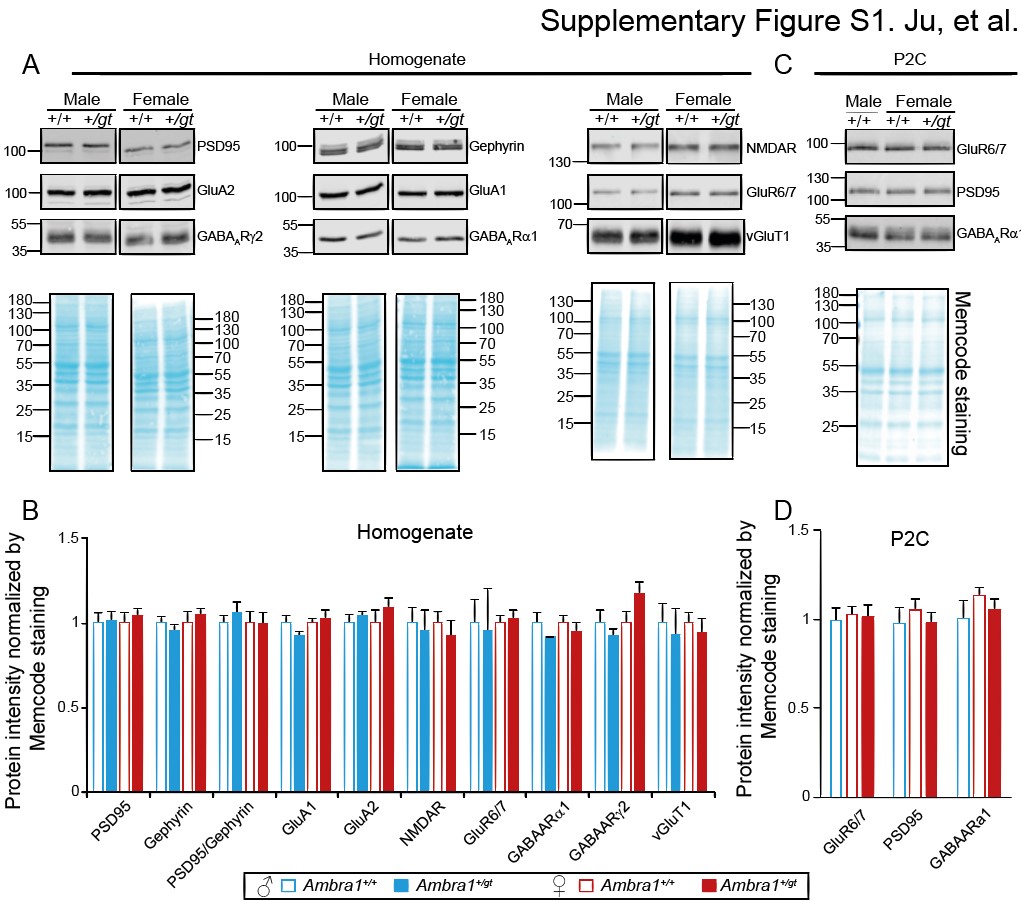

### supplementary Figure 2

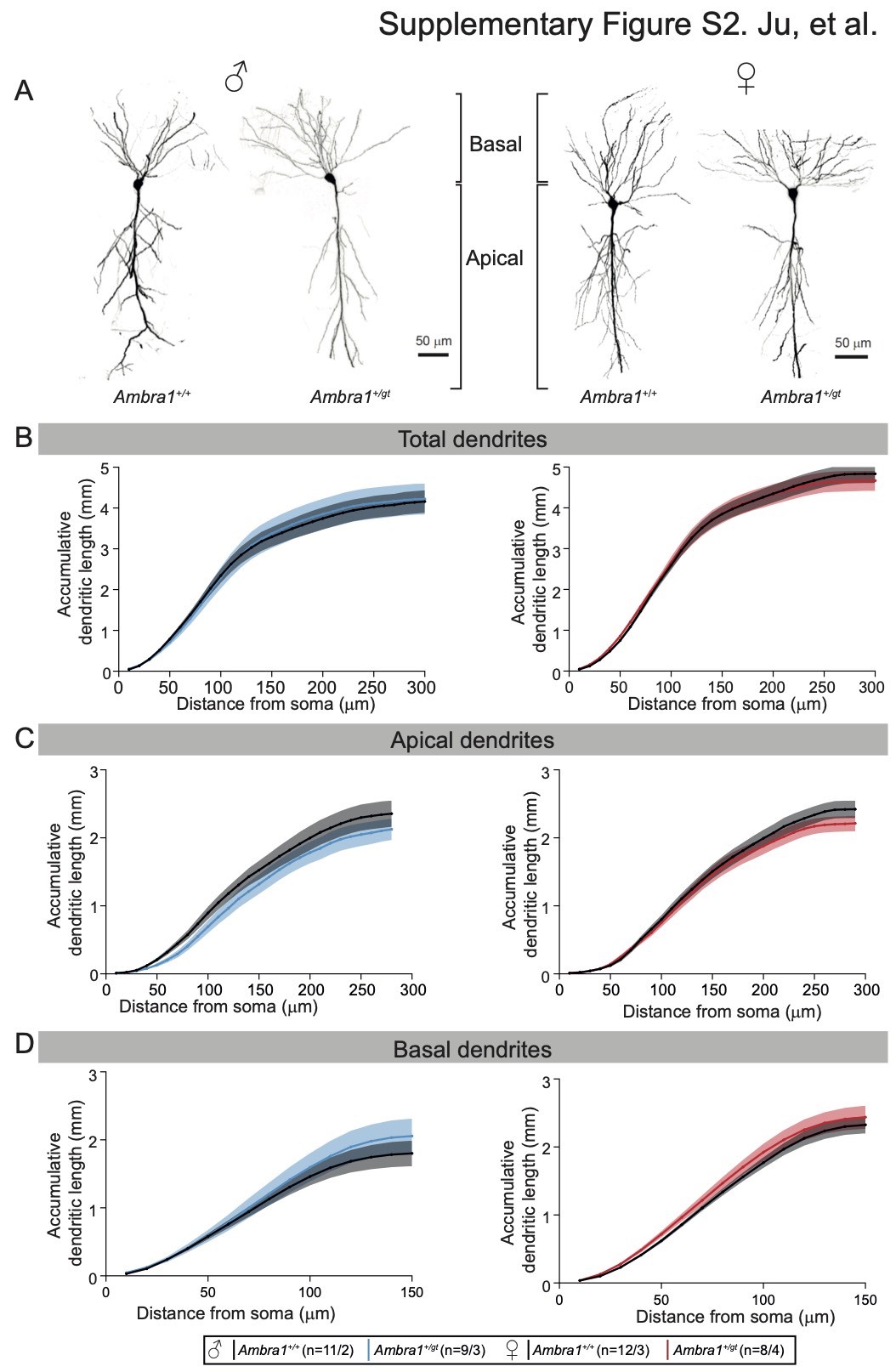

### supplementary Figure 3

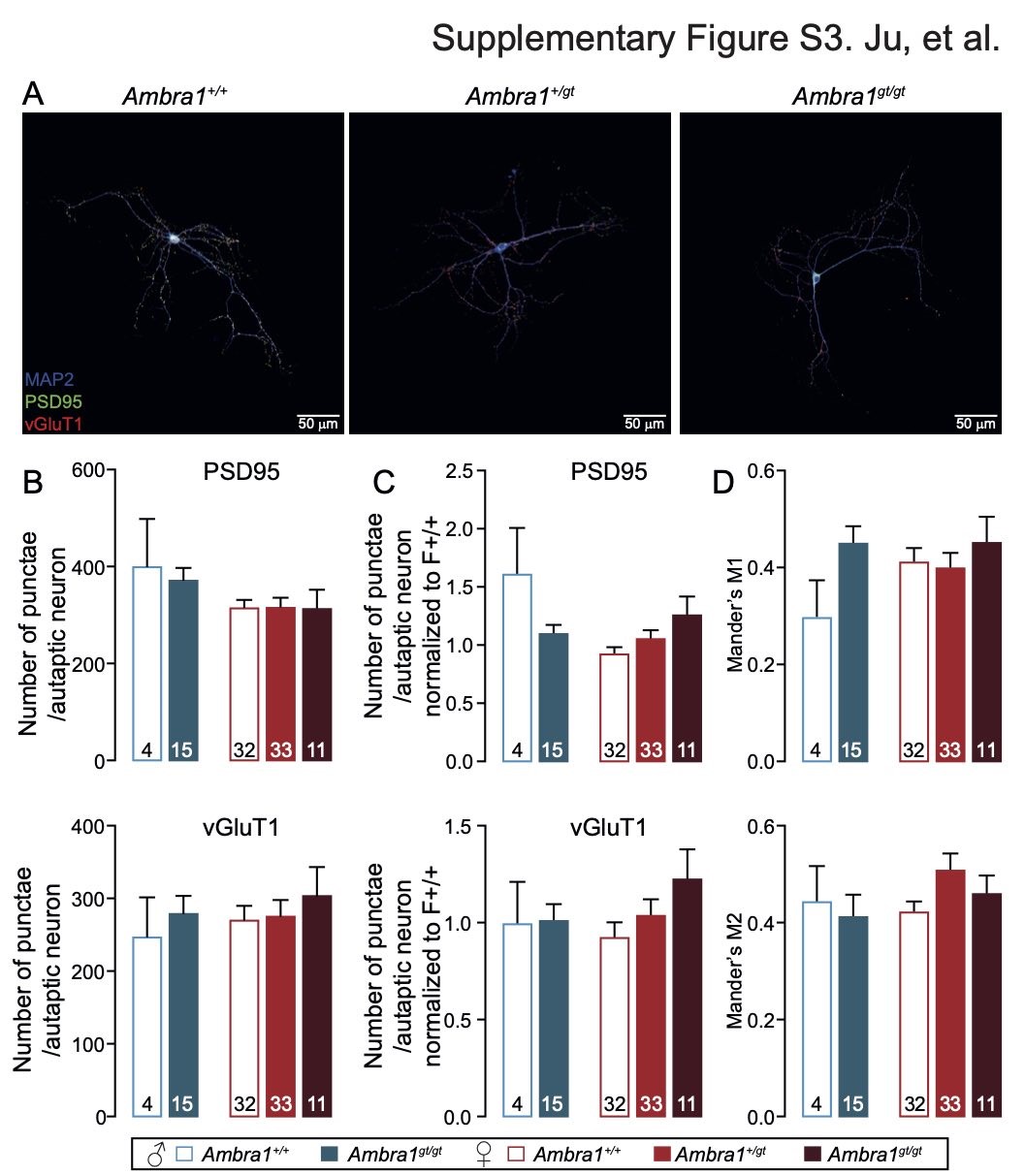
